# The Progestin DMPA is associated with increased CD 3 and CD 4 T-Lymphocytes among women living with HIV

**DOI:** 10.1101/066225

**Authors:** Edwin Walong, Anne Barasa, Christopher Gontier

## Abstract

**Introduction:** Establishment of peripheral blood CD 3 and CD 4 lymphocyte counts is useful for immunological monitoring and staging of HIV. This forms the basis of management. This study evaluates total T lymphocyte and CD4 lymphocyte counts among women on DMPA and compares this to matched controls that were not on hormonal contraception.

**Materials and Methods:** This case control study was conducted in the western Kenya city of Kisumu. Participants were WHO stage I and II ART naïve HIV-positive women. The cases were enrolled in the institutional family planning clinic and must have had an intramuscular injection of DMPA within a three month period. We used a standard provider initiated questionnaire to collect social and demographic information. Peripheral blood CD 3 and CD 4 lymphocyte counts were determined using BD–Facs-count ™. Data was entered and analysed using SPSS^®^ Version 17.

**Results:** A total of 138 participants were recruited into the study, 66 controls and 54 cases. The median ages were 26 and 28 respectively. The median CD 3 lymphocyte counts among controls and cases were 1628 and (p=0.004) while median CD4 lymphocyte counts are 649 (p=0.02).

**Conclusion:** Use of the progestin DMPA is associated with an increase in median in CD3 and CD 4 I lymphocyte counts. Although the clinical benefits of this increase is unclear, DMPA is safe for use among women living with HIV.

## Introduction

Peripheral blood T lymphocytes counts, in particular the CD 4 subset of T lymphocytes, are essential parameters for immunological monitoring of HIV/AIDS and early detection of treatment failure (1). This forms the basis for initiation of antiretroviral therapy and substitution of therapy in treatment failure (2). Variability in CD 4 lymphocyte counts can lead to inappropriate clinical decisions (3).

In the western Kenya city of Kisumu, the progestin depot medroxy-progesterone acetate (DMPA) is the most preferred contraceptive method (4). This injectable contraceptive is discrete, non toxic and effective and therefore preferred by most women in this region. Heffron et al in 2013 showed that there is no evidence that DMPA use by HIV positive women is associated with adverse clinical progress (5).

Collazos et al report lower than expected CD 4 counts among women in whom adequate viral suppression has been accomplished by HAART administration (6). It is likely that DMPA use is associated with reduction in CD4 lymphocyte counts, whose interpretation my lead to inaccurate HIV clinical staging and inappropriate clinical decisions (1). The objective of this study is to evaluate peripheral blood CD 3 and CD 4 lymphocyte counts among asymptomatic healthy ART Naive women living with HIV.

## Materials and Methods

This case control study was performed in the Family AIDS Care and Education (FACES) clinic and laboratory in Kisumu, Kenya from August 2010 to February 2011. The clinic is run by the Research Care and Training Programme (RCTP) of the Kenya Medical Research Institute (KEMRI). Participants were healthy HIV positive women, not on antiretroviral therapy (ART). Cases were women who had received at least one dose of DMPA as a deep intramuscular injection within a three month period. Controls were women who were not on hormonal contraception with regular menstrual cycles.

Informed consent was administered and a standard questionnaire was administered by the health care provider, through which social and demographic characteristics of the participants were documented.

Peripheral blood was collected from the antecubital fossa into a specimen bottle containing EDTA anticoagulant and submitted to the laboratory for CD 3 and CD 4 lymphocyte count determination.

CD 3 and CD 4 lymphocyte count determination was performed on the sample of whole blood using the automated Beckton Dickinson FACS Count^®^ according to standard operating procedure.

Data was entered into SPSS v 17. Presentations of dependent variables are in tables. Evaluation of normality using Shapiro-Wilk’s test was performed and bivariate statistical tests (Mann-Whitney) was applied to evaluate median CD 3 and CD 4 values.

## Results

A total of 138 participants were recruited, 52 cases and 66 controls. Their social and demographic data is summarised in table 1. We further report that higher median CD3 and CD4 were observed among cases as compared to cases. Cases further had greater interquartile ranges. (Table 3).

**Table 1:** SOCIAL, DEMOGRAPHIC CHARACTERISTICS AND LABORATORY ANALYSIS.

|  | Controls<br>(n=66) | Case (n=52) | Test stat. | P value |
| --- | --- | --- | --- | --- |
| **Age in years.** |  |  | Ranksum test | 0.021 |
| median(IQR) | 28 (25,34) | 26 (23,28.5) |  |  |
| **marital status (percentage)** |  |  | Fisher's exact | 0.345 |
| Single | 17 (68) | 8 (32) |  |  |
| Married | 39 (50) | 39 (50) |  |  |
| Separated | 5 (50) | 5 (50) |  |  |
| Divorced | 3 (75) | 1 (25) |  |  |
| Widowed | 6 (75) | 2 (25) |  |  |
| **highest level of education** |  |  | Fisher's exact | 0.095 |
| Primary | 38 (50.7) | 37 (49.3) |  |  |
| Secondary | 21 (56.8) | 16 (43.2) |  |  |
| College | 7 (87.5) | 1 (12.5) |  |  |
| University | 3 (100) | 0 (0) |  |  |
| adult education | 1 (100) | 0 (0) |  |  |
| **occupation of participant** |  |  | Fisher's exact | 0.271 |
| Housewife | 7 (63.6) | 4 (36.4) |  |  |
| Farmer | 4 (80) | 1 (20) |  |  |
| Unemployed | 9 (40.9) | 13 (59.1) |  |  |
| Professional | 13 (72.2) | 5 (27.8) |  |  |
| Other | 37 (54.4) | 31 (45.6) |  |  |
| **sec2.drug use** |  |  | Fisher's exact | 1 |
| Alcohol | 2 (50) | 2 (50) |  |  |
| None | 67 (56.3) | 52 (43.7) |  |  |
| **age at menarch** |  |  | Ranksum test | 0.422 |
| median(IQR) | 15 (14,16) | 14 (13.5,15) |  |  |
| **degree of blood loss** |  |  | Chisq. (2 df) = 4 | 0.127 |
| light(1pad/24hrs) | 9 (50) | 9 (50) |  |  |
| moderate(2-3pads/24hrs) | 47 (52.8) | 42 (47.2) |  |  |
| heavy(>3pads/24hrs) | 12 (80) | 3 (20) |  |  |
| **ever been pregnant** |  |  | Fisher's exact | 0.042 |
| Yes | 61 (53) | 54 (47) |  |  |
| No | 9 (90) | 1 (10) |  |  |
| **number of live births** |  |  | Ranksum test | 0.195 |
| median(IQR) | 2 (1,3) | 2 (2,3) |  |  |
| number of still births |  |  | Ranksum test | 0.232 |
| median(IQR) | 0 (0,0) | 0 (0,0) |  |  |
| number of abortions |  |  | Ranksum test | 0.393 |
| median(IQR) | 0 (0,0) | 0 (0,0) |  |  |
| current fp method |  |  | Fisher's exact | < 0.001 |
| Natural | 4 (100) | 0 (0) |  |  |
| male/female condoms | 4 (100) | 0 (0) |  |  |
| Injectable | 0 (0) | 18 (100) |  |  |
| condoms & injectable | 0 (0) | 37 (100) |  |  |
| fp method last 6months |  |  | Fisher's exact | < 0.001 |
| male/female condoms | 11 (91.7) | 1 (8.3) |  |  |
| Injectable | 4 (21.1) | 15 (78.9) |  |  |
| condoms & injectables | 1 (2.8) | 35 (97.2) |  |  |
| hvs for gc culture |  |  | Fisher's exact | 1 |
| Positive | 1 (100) | 0 (0) |  |  |
| Negative | 69 (55.6) | 55 (44.4) |  |  |
| microscopy for candida |  |  | Fisher's exact | 1 |
| Positive | 4 (57.1) | 3 (42.9) |  |  |
| microscopy for bv |  |  | Chi Square | 0.979 |
| Positive | 15 (57.7) | 11 (42.3) |  |  |
| Negative | 55 (55.6) | 44 (44.4) |  |  |
| microscopy for tvaginalis |  |  | Fisher's exact | 1 |
| Positive | 1 (100) | 0 (0) |  |  |
| Negative | 69 (55.6) | 55 (44.4) |  |  |
| cervical smear for papanicolau staining |  |  | Fisher's exact | 0.005 |
| Normal | 67 (61.5) | 42 (38.5) |  |  |
| Reactive | 2 (22.2) | 7 (77.8) |  |  |
| low grade | 0 (0) | 4 (100) |  |  |
| high grade | 1 (33.3) | 2 (66.7) |  |  |

**Table 2:** PERIPHERAL BLOOD CD3 AND CD4 LYMPHOCYTE COUNTS AMONG STUDY PARTICIPANTS

| Variable | Control | Case | Test Statistic | Pvalue |
| --- | --- | --- | --- | --- |
| CD 4 lymphocyte counts (per microliter)<br>median(IQR) | 573.5 (474,726.5) | 649 (524,916.5) | Ranksum test | 0.02 |
| Cd 3 lymphocyte counts (per microliter)<br>median(IQR) | 1603 (1271,2006) | 1935 (1628,2294) | Ranksum test | 0.004 |

This table shows the median lymphocyte counts among cases and controls. Higher median values are observed among women on DMPA

## Discussion

This study found an increase in median CD3 and CD 4 lymphocyte counts among women on DMPA. These are unexpected as in vitro studies on cell lines as well as animal studies show reduced T lymphocyte counts (7). In addition, longitudinal studies on large cohorts of women in Kenyan and Ugandan sites did not show suppression of CD 4 counts (5).

This study specifically selected a population of ART naïve individuals whose CD 4 counts were above 350 per microliter, the threshold by which current guidelines indicate initiation of antiretroviral therapy. In light of these, laboratory reference ranges among women on DMPA should be established (9).

While it would be expected that women with lower CD 4 counts due to DMPA use would have an increased risk of progression to clinical stages 3 and 4, there was no evidence of increase in mucosal or other infections among these women or other clinical adverse effects. There are no reported clinical benefits of DMPA use due to elevated lymphocyte counts (5)(7). One likely benefit is the effect of DMPA also inhibits helminthic and protozoal infections, whose prevalence in Western Kenya is high and may have contributed to higher lymphocyte counts (7). Effective contraception with reduction of chances of pregnancy leads to overall improvement in the health of women (4).

While this study did not evaluate the clinical significance of these changes, we postulate that the reported increase in CD3 and CD 4 lymphocyte counts due to DMPA use may lead to inaccurate interpretation of immunological monitoring of HIV (1). This may necessitate evaluation of reference ranges among these participants and correlation with systemic HIV 1 viral loads. There is a high risk of misclassification when immunological monitoring is applied in women on DMPA.

## Conclusion

Use of the Progestin DMPA is associated with higher median CD 3 and CD 4 counts. This may be as a result of more favourable health care indices associated with contraceptive use among women living with HIV. The impact of these changes upon immunological monitoring of HIV is unclear. This should be further evaluated preferably using longitudinal studies.

## Author Contributions

EW designed the study, recruited the participants, performed the CD 3 and CD 4 assays and wrote the manuscript. AB and CG participated in the design of the study, performed data analysis and participated in the writing of the manuscript. All authors read and approved the final manuscript

## Acknowledgements

The clients and staff of the Family Aids Care and Education (FACES) clinic, Research Care and Training Programme (RCTP), Center for Microbiology Research (CMR), Kenya Medical Research Institute (KEMRI).

## List of Abbreviations

AIDS: Acquired Immune Deficiency Syndrome
ART: Antiretroviral Therapy
CD: Cluster of Differentiation
CMR: Center for Microbiology Research
DMPA: Depot Medroxyprogesterone Acetate
EDTA: Ethylenediaminetetracetic Acid
FACS: Florescent Activated Cell Sorter
FACES: Family Aids and Education Services
KEMRI: Kenya Medical Research Institute
RCTP: Research Care and Training Program
WHO: World Health Organisation

